# The characteristics of the HIV-1 Env glycoprotein contribute to viral pathogenesis

**DOI:** 10.1101/2021.07.07.451566

**Authors:** Silvia Pérez-Yanes, Maria Pernas, Silvia Marfil, Romina Cabrera-Rodríguez, Raquel Ortiz, Carla Rovirosa, Judith Estévez-Herrera, Isabel Olivares, Concepción Casado, Cecilio Lopez-Galindez, Julià Blanco, Agustin Valenzuela-Fernández

**Affiliations:** Laboratorio de Inmunología Celular y Viral, Unidad de Farmacología, Sección de Medicina, Facultad de Ciencias de la Salud de la Universidad de La Laguna (ULL), Campus de Ofra s/n, 38071 Tenerife, Spain.; Unidad de Virologia Molecular, Laboratorio de Referencia e Investigación en Retrovirus. Centro Nacional de Microbiologia. Instituto de Salud Carlos III. Majadahonda 28220 Madrid. Spain.; Institut de Recerca de la Sida IrsiCaixa, Institut d’Investigació en Ciències de la Salut Germans Trias i Pujol (IGTP), 08916 Badalona, Spain; Universitat de Vic, Universitat Central de Catalunya, UVIC-UCC, 08500 Vic, Spain.

## Abstract

The understanding of HIV-1 pathogenesis and clinical progression is incomplete because of the variable contribution of host, immune and viral factors. The involvement of viral factors has been investigated in extreme clinical phenotypes from rapid progressors to long-term non-progressors (LTNPs). Among HIV-1 proteins, the envelope glycoprotein complex (Env) has concentrated many studies for its important role in the immune response and in the first steps of viral replication. In this study, we analyzed the contribution of 41 Envs from 24 patients with different clinical progression rates and viral loads (VLs), LTNP-Elite Controllers (LTNP-ECs); Viremic LTNPs (vLTNPs), and non-controller’s individuals contemporary to LTNPs or recent, named Old and Modern progressors. We analyzed the Env expression, the fusion and cell-to-cell transfer capacities as well as viral infectivity. The sequence and phylogenetic analysis of Envs were also performed. In every functional characteristic, the Envs from subjects with viral control (LTNP-ECs and vLTNPs) showed significant lower performance compared to those from the progressor individuals (Old and Modern). Regarding sequence analysis, the variable loops of the gp120 subunit of the Env (i.e., V2, V4 and mainly V5) of the progressor individuals showed longer and more glycosylated sequences than controller subjects. Therefore, HIV-1 Envs presenting poor viral functions and shorter sequences were associated with viremic control and the non-progressor clinical phenotype, whereas functional Envs were associated with the lack of virological control and progressor clinical phenotypes. These correlations support the central role of Env genotypic and phenotypic characteristics in the *in vivo* HIV-1 infection and pathogenesis.

**IMPORTANCE:** The role of the virus in the pathogenesis of HIV-1 infection has not been investigated in isolates from individuals with different progression rates. In this work, we studied the properties of the envelope glycoprotein complex (Env) in individuals with different progression rates to elucidate its role in pathogenesis. We estimated the Env expression, the CD4 binding, the fusion and cell-to-cell viral transfer capacities that affect the infectivity of the viral Envs in recombinant viruses. The Envs from individuals which control viral replication and lack clinical progression (LTNP-ECs and vLTNPs) showed lower functional capacities than from subjects with clinical progression (Old and Modern). The functional increase of the Envs characteristics was associated with an increase in viral infectivity and in increased length of variable loops and the number of glycosylation sites of the Env (gp120/SU). These results support the concept that viral characteristics contribute to viral infection and pathogenesis.

## Introduction

Pathogenesis of viral infections is the result of complex interactions between host genetics, immune responses and viral factors. In human immunodeficiency virus tye 1 (HIV-1) infection and pathogenesis, the role of host (1–6), immune (6–15) and viral factors (16–20) has been widely investigated. The interactions of these factors have been primarily studied in extreme clinical phenotypes like rapid progressors (RPs) (21, 22) or long-term non-progressors (LTNPs), LTNP-Elite Controllers (LTNP-ECs), HIV controllers or Elite suppressors (ES) (17-19, 23, 24).

Due to these entangled interactions, the investigation of the role of viral proteins and their specific properties in HIV-1 pathogenesis is challenging. Among the viral proteins, the envelope glycoprotein complex (Env) has attracted numerous studies because its essential role in the immune response and in the initial events of the HIV-1 biological cycle (25–29), i.e the binding to the cellular receptors (29–42). The binding efficiency of the viral Env to the CD4 receptor determines further steps of the viral cycle: virus-cell signaling, fusion and cell-to-cell virus transfer capabilities (18, 19, 43). HIV-1 Envs unable to stabilize microtubules (i.e., increasing post-transductional acetylation of Lys^40^ residue in *α*-tubulin), to reorganize F-actin for the delineation of pseudopod-entry virus hot zones present low CD4 binding, restricted fusion and low early infection (18, 19, 43–45).

There are few reports investigating the characteristics of viral Envs from HIV individuals with different clinical characteristics. Lassen *et al.* studied the entry efficiency of viral Envs from ES individuals relative to chronically infected viremic and chronic progressors. Envs from ES showed decreased entry efficacy and slower entry kinetics than those of chronic progressors (20). Our group studied the CD4 binding, signaling capacity, fusogenicity of viral Envs from viremic non-progressors (VNPs) that were similar to those of progressors individuals (19). In previous reports, deficient viral Env glycoproteins, because of poor CD4 binding, low transfer and signaling capacity (18) were identified in a cluster of poor replicating viruses from a group of LTNP-ECs without clinical progression for more than 20 years (17, 18). Thus, these works have stablished that viral Env play an important role in the pathogenesis control in LTNPs (17-20, 46, 47).

To further investigate the role of viral Env in HIV-1 infection and pathogenesis, in this work, we expanded our previous studies to viral Envs from other sets of LTNP-ECs and Viremic LTNPs (vLTNPs) in comparison with groups of chronic progressors. Clonal full-length *env* genes derived from viruses of individuals in these distinct clinical groups were analyzed for expression, CD4 dependent-Env-mediated fusion, cell-to-cell viral transfer and infection efficiency. This analysis permitted the establishment of a relationship between the initial events of the viral replication cycle, mediated by the viral Env characteristics, with the VL control and the clinical outcome and pathogenesis of the HIV-1 infection.

## Results

### Analysis of the characteristics of viral envelopes of viruses from different risk groups

For the investigation of the potential role of the HIV-1 Env in virological control and pathogenesis, we studied the phenotypic characteristics of 41 Envs from 24 individuals without antiviral therapy and different VLs (**Table 1**). We analyzed 10 Envs from 6 LTNP-EC individuals with undetectable VL and infected in the late 80’s and 90’s; 10 viral clones from 6 Viremic LTNPs (vLTNPs) with VL <10,000 viral copies/mL and infected in the 90’s. To ascertain that the characteristics of the Envs from these LTNPs were not due to the sampling time, we compared them with 10 Envs obtained from 6 HIV-1 individuals also infected in the same period (90’s), but with high VL>10^5^ viral copies/mL and chronic infection; these Env were designated Old. Finally, we studied 11 viral clones from 6 chronic individuals infected between 2013-2014 with VL>10^4^ viral copies/mL and named Modern. The main characteristics of the participants are summarized in **Table 1**.

**Table 1.**
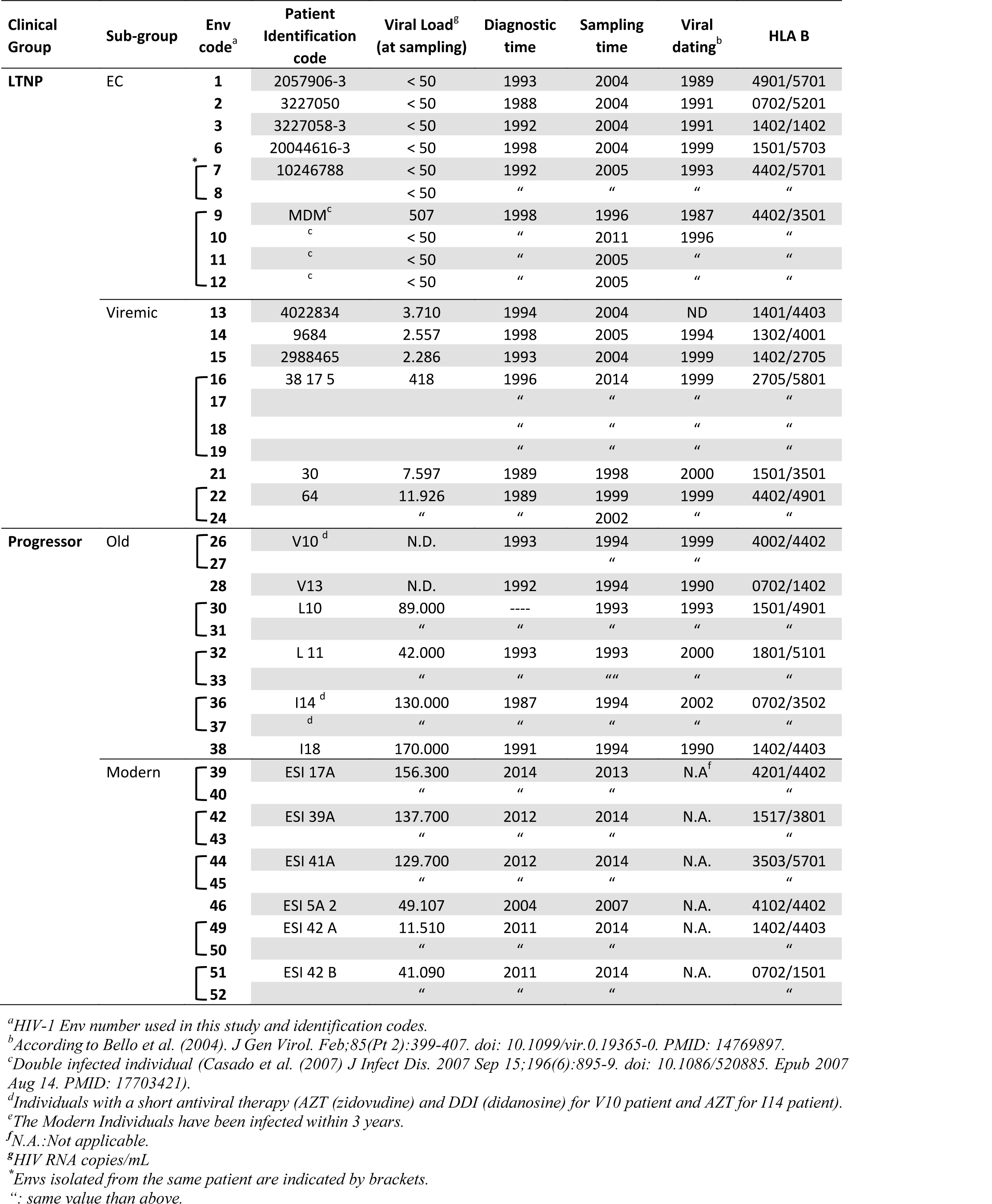
Epidemiological, clinical and host characteristics of the viral Envs.

We first analyzed the potential differences in the expression between the Env clones from the clinical groups, by measuring their cell-surface expression levels in HEK-293T cells (**Figure 1A***, shows study scheme*, and **Figure 2**). Although we observed a progressive augmentation of Env expression in viral clones derived from patients that do not control viremia (i.e., Old and Modern patients) compared to LTNPs (EC and Viremic), this increase did not reach statistical significance (**Figure 2**). Thus, the expression capability of the viral Envs appears to not contribute to the differences in VL and pathogenesis between groups.

**Figure 1.**
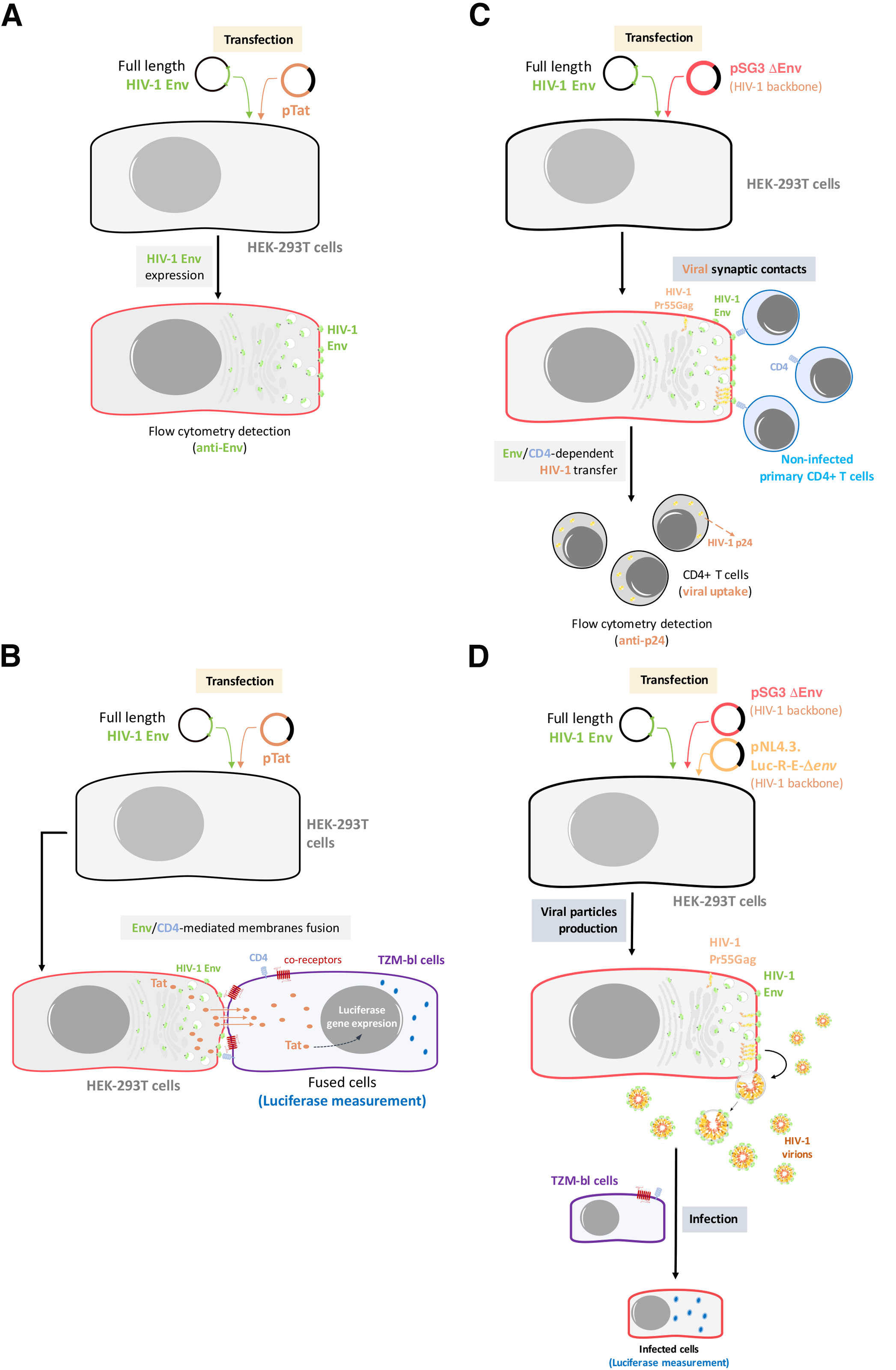
Outline of the experimental model used for the analysis of Env expression, Env-mediated cell-to-cell fusion, viral transfer and viral infectivity. (A) Env expression: HEK-293T cells will be co-transfected with primary of reference full-length viral *env* and a ptat Δ*env* HIV-1 expression plasmid, allowing Env cell-surface expression in a viral production context. Cell-surface Env expression will be then analyzed by flow cytometry using specific anti-Env antibody. (B) Env-mediated fusion activity: after 24 hours, effector HEK-293T cells producing HIV-1 particles bearing primary or reference Envs will be co-cultured with TZM-bl cells to force synapsis formation and CD4-mediated binding of budding particles to target cells. (C) Env-mediated viral transfer: HEK-293T cells producing HIV-1 particles carrying primary or reference Envs will be co-cultured with primary CD4+ T cells. Then, HIV-1 transfer will be analyzed by flow cytometry using specific anti-p24 antibody in target CD4+ T cells. (D) Env-mediated viral infection: TZM-bl cells will be infected with serial dilutions of viral particles obtained from transfected HEK-293T and carrying the different primary or reference HIV-1 Envs. After 48 hours, infectivity capacity will be analyzed by quantifying luciferase assay in infected TZM-bl cells.

**Figure 2.**
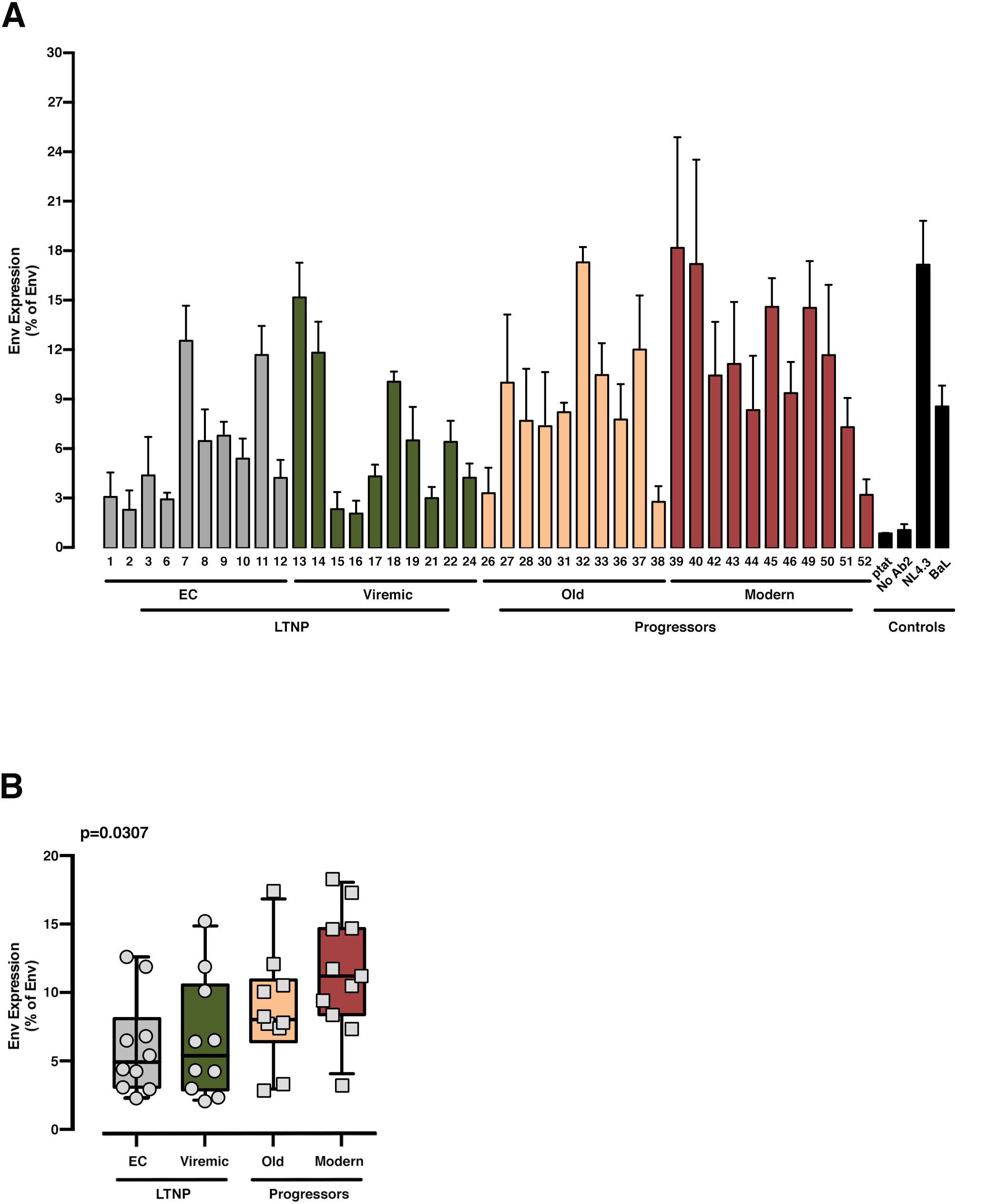
Analysis of the expression of the different HIV-1-Env glycoproteins from LTNP-EC, Viremic LTNP and control progressors patients. Flow cytometry analysis of the cell-surface expression level of the assayed HIV-1 Envs in HEK-293T cells from LTNP-EC (*gray bars*), vLTNP (*green bars*), Old (*orange bars*) and Modern individuals (*red bars*) or reference HIV-1 viral strains (ptat, No Ab2, NL4.3 and BaL, *black bars*). Env protein expression for each patient (A) and Env protein expression in each group of patients comparing mean values between each group (Kruskal-Wallis, Dunn’s Multiple Comparisons Test) (B); p value for comparison between all groups is shown, *top left*. Values are mean ± S.E.M. of three independent experiments.

### Analysis of cell-to-cell membrane fusion and viral transfer capacity of viral envelopes

A key process for HIV Env-mediated infection is the interaction of the Env complex with the CD4 receptor. When this interaction is functionally efficient, viral transfer through synaptic contacts or fusion pore formation are triggered during cell-to-cell or virus-to-cell contacts, repectively (18, 19, 43, 45, 48). We examined the viral Env/CD4 interaction and the efficiency of subsequent functions, measuring the membrane fusion capacity of the Envs (**Figure 1B**, *shows study scheme*) in co-cultures between Env-expressing HEK-293T and HIV-permissive target TZM-bl cells (**Figure 3**). To fully characterize our experimental models, we used the Envs from reference HIV-1_BaL_ (CCR5-tropic) and HIV-1_NL4.3_ (CXCR4-tropic) viruses (**Figure 3** and **4**). This fusion assay yielded lower fusion values for Envs of viruses from LTNP-ECs and from vLTNPs than for Old and Modern progressors, and attaining statistical significance between LTNPs (EC and Viremic) and Modern Envs glycoproteins (**Figure 3B**).

**Figure 3.**
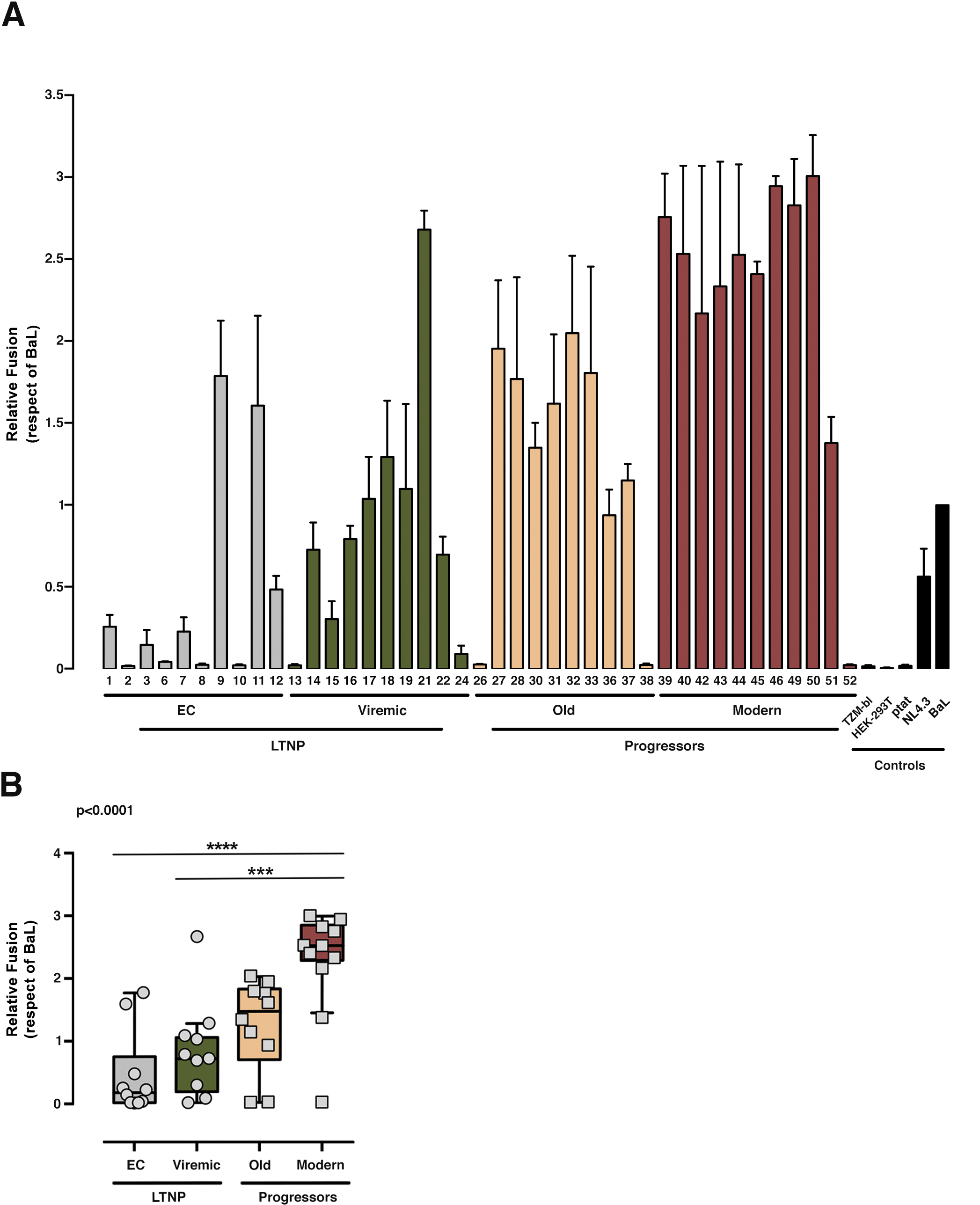
Analysis of membranes fusion-phenotypic features of HIV-1 Envs isolated from LTNP-EC, viremic LTNP and P individuals. Analysis of the ability to induce cell-to-cell fusion of HIV-1 Env proteins obtained from LTNP-EC (*gray bars*), vLTNP (*green bars*), Old (*orange bars*) and Modern individuals (*red bars*) or reference HIV-1 viral strains (ptat, NL4.3 and BaL, *black bars*). (A) Env fusogenic activity for each patient in each group. (B) Relative fusion activity of the full Env collection compared to the BaL control established at 100% and grouped in the different groups of patients. Values are mean ± S.E.M. of three independent experiments. Statistical analysis was performed using Kruskal-Wallis, Dunn’s Multiple Comparisons Test; p value for comparison between all groups is shown, *top left*.

**Figure 4.**
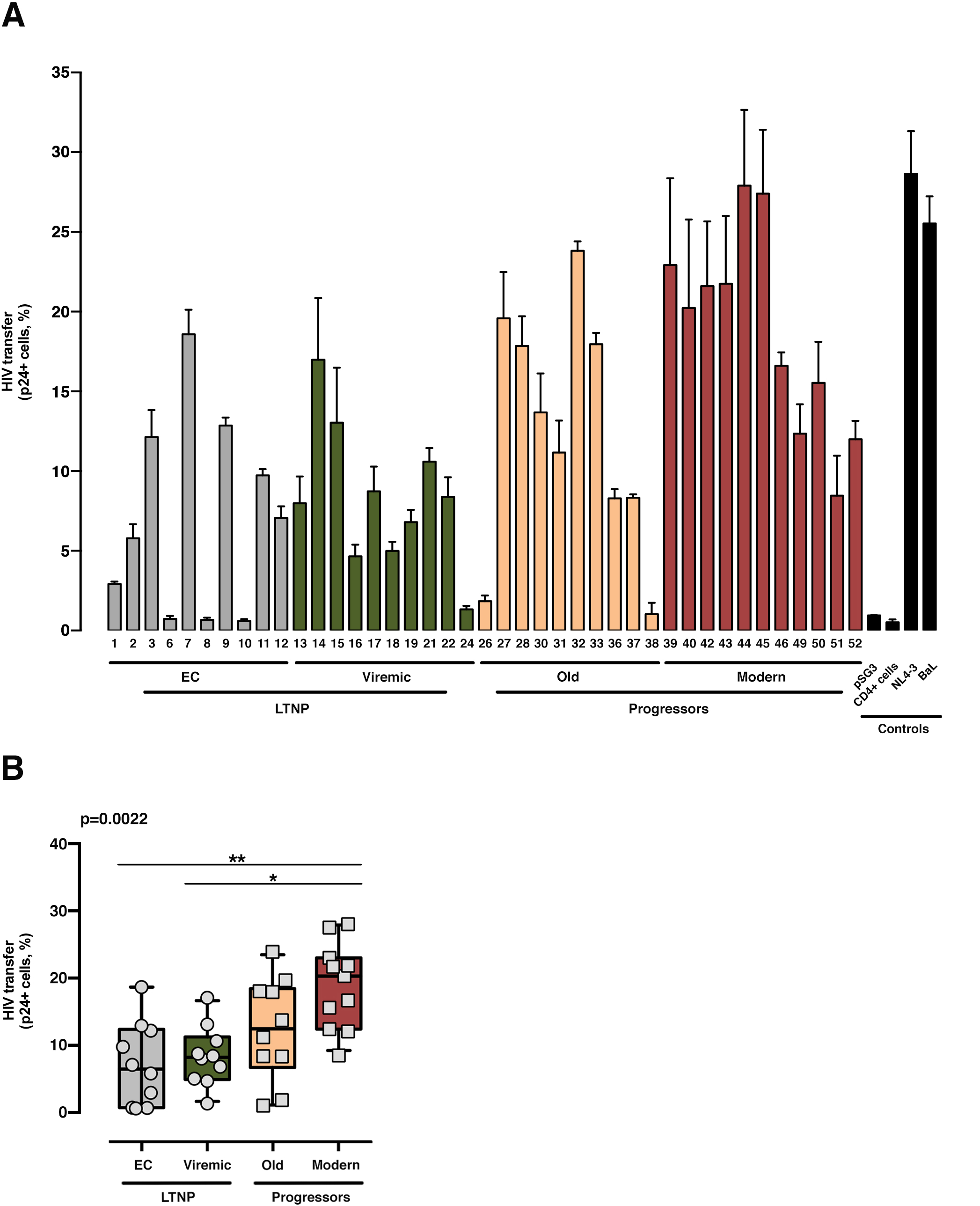
Analysis of HIV-1 Env-mediated cell-to-cell viral transfer. Analysis of the ability to induce cell-to-cell virus transfer of HIV-1 Env proteins obtained from LTNP-EC (*gray bars*), vLTNP (*green bars*), Old Patients (*orange bars*), recent patients (Moderns) (*red bars*) or reference HIV-1 viral strains (pSG3, CD4+ cells, NL4.3 and BaL, *black bars*). Analysis of HIV-1 Env-mediated cell-to-cell viral transfer for each patient (A) and in each group where P values compare medians between groups using a nonparametric Kruskal-Wallis Test (Kruskal-Wallis, Dunn’s Multiple Comparisons Test) (B); p value for comparison between all groups is shown, *top left*. Values are mean ± S.E.M. of two independent experiments.

Next, we assayed the CD4-dependent cell-to-cell virus transfer capacity of the viral envelopes. This experiment was performed co-culturing Env-expressing HEK-293T cells with unstimulated primary CD4+ T lymphocytes as target cells (**Figure 1C**, *shows study scheme*, and Materials and methods). In this assay, we forced the formation of virological synapses between virus-effector HEK-293T cells expressing the different Envs together with the structural HIV Gag polyprotein, and fresh primary CD4+ T cells from healthy donors (**Figure 1C**, *shows study scheme*). The Envs from the LTNPs (EC and Viremic) individuals displayed a lower ability to transfer viral particles to primary CD4+ T lymphocytes than Envs from Old individuals and significantly lower than from Modern participants (p<0.0022 between all groups) (**Figure 4**). These data suggest that the Envs from LTNP-EC viruses had an impaired binding to the cell-surface CD4 receptor and that this impairment was progressively overcome in the Envs from individuals from the other groups with less control of viral replication, and higher VL.

Thus, the phenotypic characterization of the Envs of viruses from subjects with distinct progression rates confirmed that LTNP-ECs and vLTNPs presented viruses with an impaired Env CD4-associated functions and a significant lower fusogenic and transfer capacity, in comparison with viruses from the viremic groups: These lower characteristics were also linked with the low VL detected in these subjects (**Figures 3** and **4**). We also observed a functional improvement in the viral Envs from the LTNP-EC and vLTNP individuals to those of chronic Modern glycoproteins: These data support that the deficient Env fusion and transfer capacities observed in the Envs of viruses from LTNP-EC and vLTNP phenotypes have been enhanced in the viruses from individuals with progressive infection, particularly in those of the Modern group.

### Infectivity of recombinant viruses with the analyzed envelopes

For the exploration of the potential consequences of these Env properties in virus biology, we estimated the infectivity of recombinant viruses bearing the Env from the different HIV+ phenotypic groups in TZM-bl cells (**Figure 5** and **Figure 1D**, *shows study scheme*). Viral Envs from the LTNP-EC group showed the lowest infectivity values, whereas the Modern Envs produced the higher titers. The viruses from vLTNPs displayed higher titers than LTNP-ECs but lower than those from Old individuals. Recombinant viruses from individuals with high VL and progressive infection (Old and Modern) have higher infectivity rates than those with viral control (EC and Viremic). These results explain why the viral properties analyzed (binding, fusion and transfer) have a significant impact in viral infectivity with an important effect in the biology of HIV-1 and viral pathogenesis.

**Figure 5.**
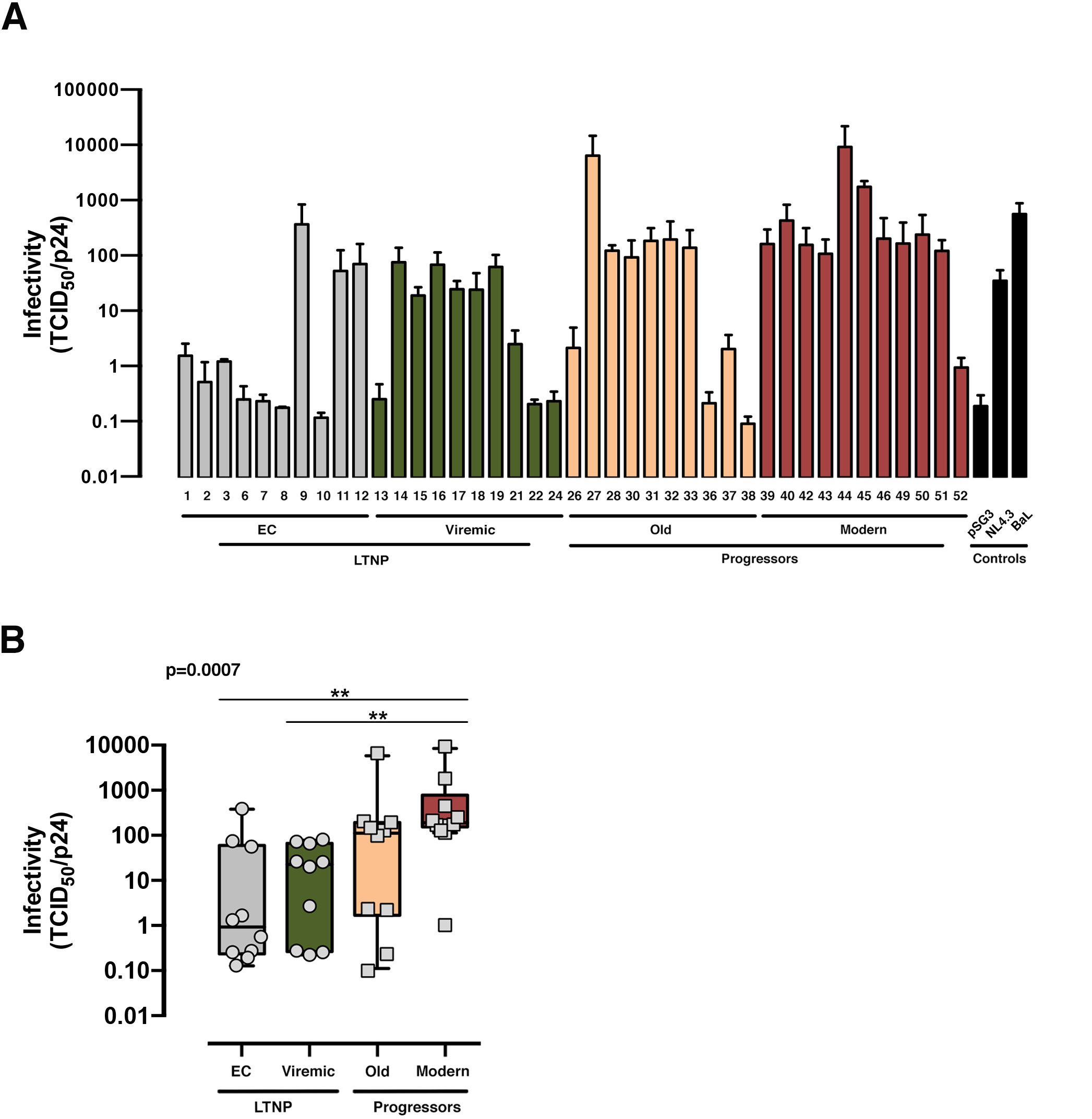
Viral infectivity of the viral Envs. Analysis of the infecivity (TCID_50_ value normalized by viral p24 input) of the different of HIV-1 Env proteins obtained from LTNP-EC (*gray bars*), vLTNP (*green bars*), Old (*orange bars*) and Moderns (*red bars*) patients or reference HIV-1 viral strains (pSG3, NL4.3 and BaL, *black bars*). Analysis of Env infectivity for each patient (A) and in each group where P values compare medians between groups using a nonparametric Kruskal-Wallis, Dunn’s Multiple Comparisons Test (B); p value for comparison between all groups is shown, *top left*. Values are mean ± S.E.M. of three independent experiments.

### Correlation between viral characteristics of the envelopes

A significant correlation was observed between the HIV-1 Env-triggered cell-to-cell transfer data, which is directly mediated by Env/CD4 binding, with Env-mediated infectivity and fusogenicity (**Figure 6**). In all viral characteristics, the Envs from subjects with virological control (EC and Viremic) showed the lower values, whereas those from the non-controlling individuals (Old and Modern) had the higher values. Therefore, HIV-1 Envs displaying poor viral functions, because of the poor binding of the viral Env to the CD4, correlated with viremic control and non-progressor clinical phenotypes. In contrast, functional Envs are associated with the lack of viremic control and the progressor clinical phenotypes. These statistical correlations support the role of viral properties in the viral phenotype that contributes to HIV-1 infection, disease progression and pathogenesis.

**Figure 6.**
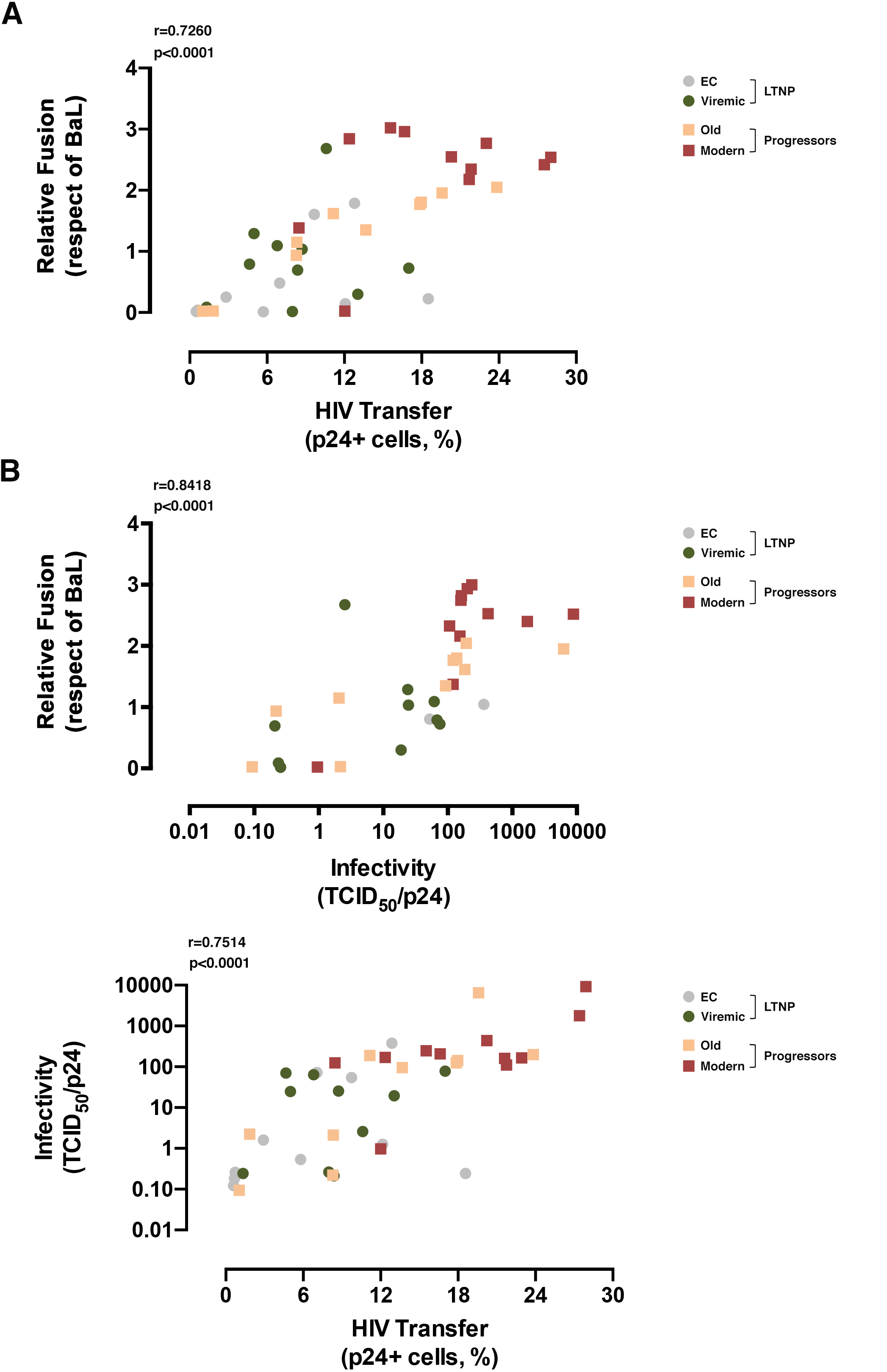
Analysis of the correlation of the fusion, transfer and viral infectivity Env characteristics between groups. (A) Correlation between Relative fusion and HIV Transfer of all Envs of the different groups LTNP-EC (*gray circle*), vLTNP (*green circle*), Old patients (*orange square*) and Modern patients (*red square*). The correlation was calculated with a nonparametric Spearman test. (B) Correlation between Relative fusion and Infectivity (TCID_50_ value normalized by viral p24 input) of all Envs of the different groups LTNP-EC (*gray circle*), vLTNP (*green circle*), Old patients (*orange square*) and Modern patients (*red square*). The correlation was calculated with a nonparametric Spearman test. (C) Correlation between Infectivty and HIV Transfer of all Envs of the differents groups LTNP-EC (*gray circle*), vLTNP (*green circle*), Old patients (*orange square*) and recent patients Moderns) (*red square*) is shown. The correlation was calculated with a nonparametric Spearman test. Values are mean ± S.E.M. of three independent experiments; p value for comparison between all groups is shown, *top left*.

### Analysis of the viral envelope sequences

For the search of potential mechanisms involved in the changes of the characteristics among the different Envs sets, we analyzed the Env amino-acid (aa) sequences that could be associated with the distinct clinical phenotypes. Initially, we performed a phylogenetic reconstruction from *env* aa sequences together with other aa sequences obtained from HIV-1 Spanish individuals. All aa sequences analized correspond to HIV-1 subtype B. This analysis did not reveal phylogenetic relationships between the different groups analysed and no clustering except for those aa sequences obtained from the same individual (**Figure 7**). Envs from LTNP-ECs and one vLTNPs grouped in short branches, as a consequence of the viral and evolutionary control, whereas long branch length was observed in the sequences obtained from non-controller patients (Old and Modern), because of the higher replication and viral evolution in these individuals.

**Figure 7.**
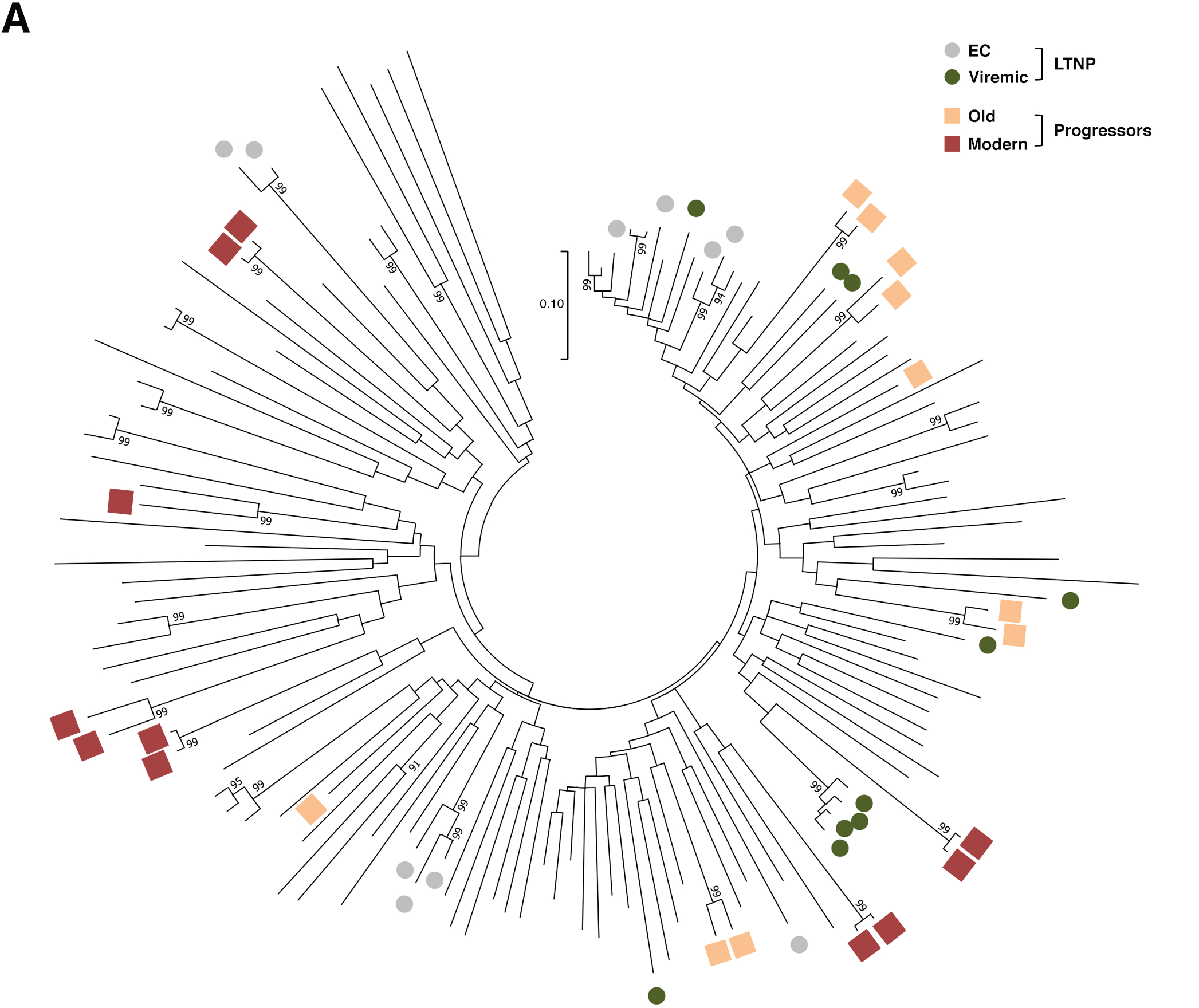
Phylogenetic analysis of the vial Envs. The evolutionary history of the Env aa sequences was inferred as described in Materials and Methods using the Maximum Likelihood method and JTT matrix-based model (109). The tree with the highest log likelihood (-49687,86) is shown. The percentage of trees in which the associated taxa clustered together is shown next to the branches. Evolutionary analyses were conducted in MEGA X (110).

We then carried out a comprehensive study of the protein sequences focusing in the variable loops and their associated potential N-linked glycosylation sites (PNGs) in the gp120 subunit of the Env. In general, as previously reported, there is a trend in the HIV-1 viral Env to gain length and glycosylation sites along the epidemic (49–51). This increasing trend is also found in our work where viruses from the LTNPs (EC, Viremic) and Old Envs isolated in the 90’s showed shorter lengths than those of the Modern group obtained in 2013-2014 (**Table 2**). The V3 loop was the most conserved and constant region in length and glycosylation sites (**Table 2** and **Figure 8**), while the other loops showed length increases predominantly in the V2 and V5 loops that were reproduced in the total length (**Table 2** and **Figure 8**). The only statistical differences were noticed between the total length in the LTNPs (EC and Viremic) versus Old and Modern Envs in the V2 and V5 regions (**Figure 8**).

**Figure 8.**
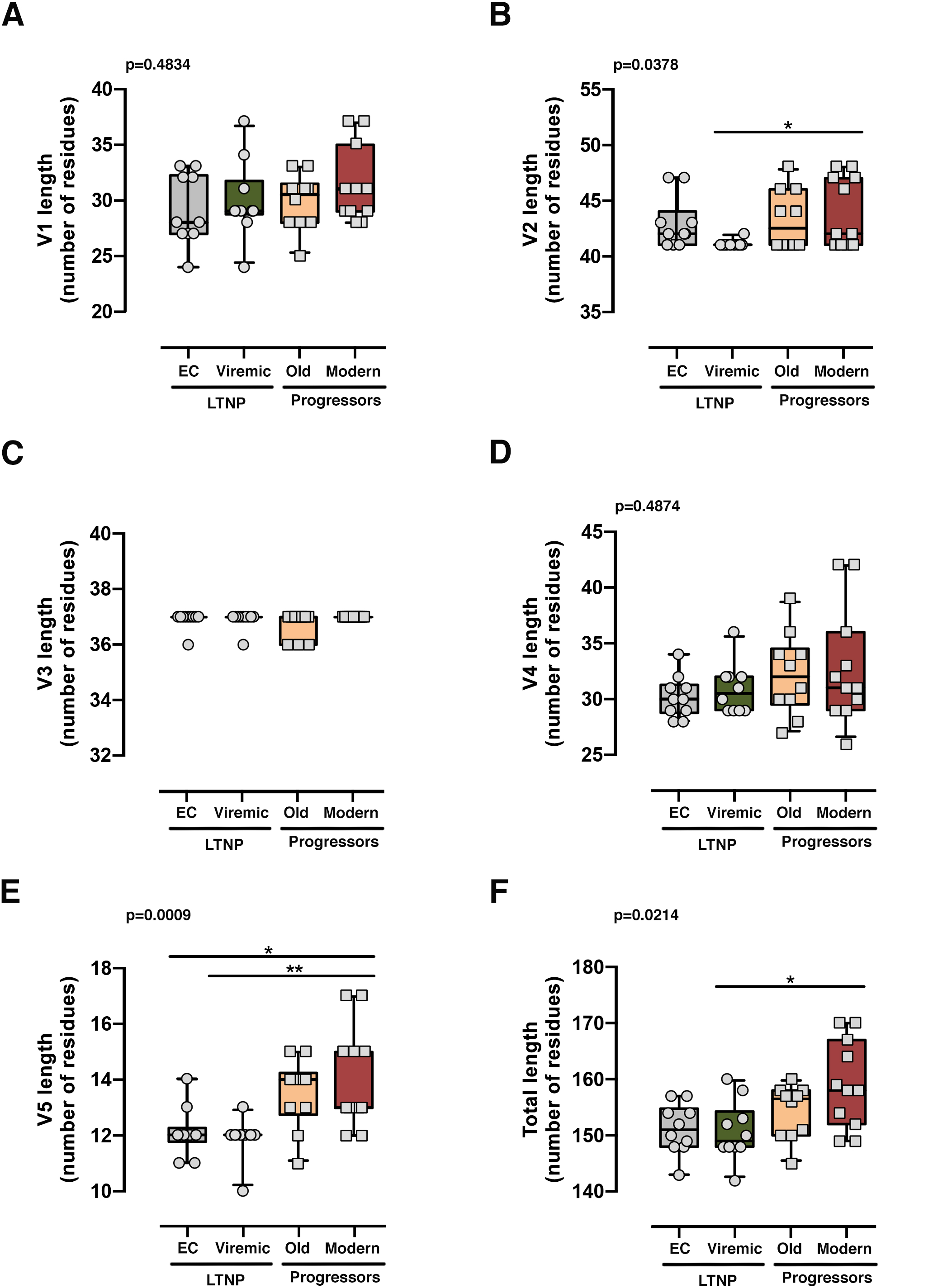
Analysis of the length and glycosylation sites in the loops of the Envs from the different groups. Analysis of the length of each variable loops V1 (A), V2 (B), V3 (C), V4 (D), V5 (E) and all variable loops together (F). The results were grouped (LTNP-ECs: *gray bar*, vLTNPs: *green bar*, Old patients: *orange bar*, and recent patients (Moderns): *red bar*) and compared using a nonparametric Kruskal-Wallis, Dunn’s Multiple Comparisons Test; p value for comparison between all groups is shown, *top left*. Values are mean ± S.E.M. of three independent experiments.

**Table 2.**
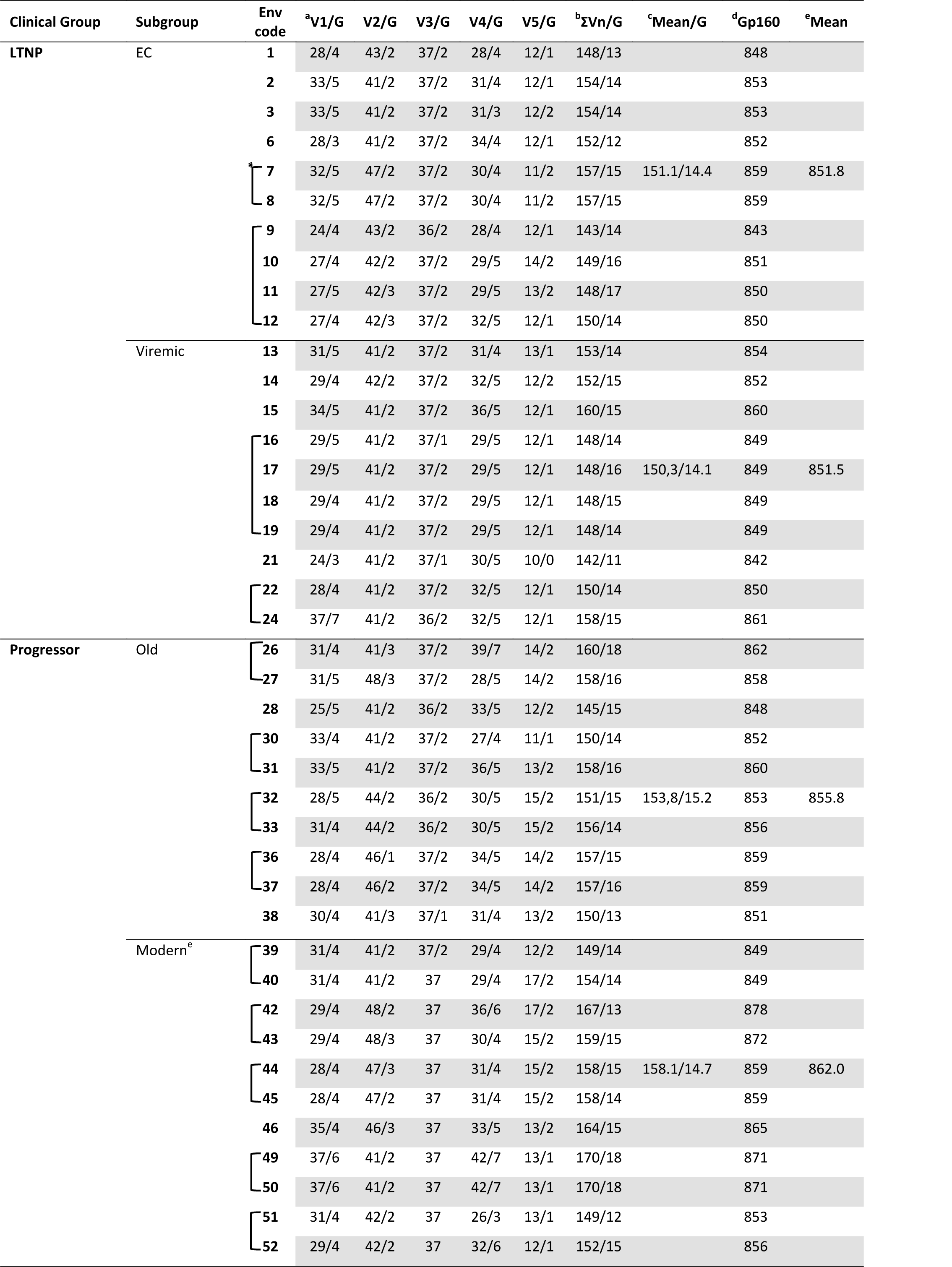
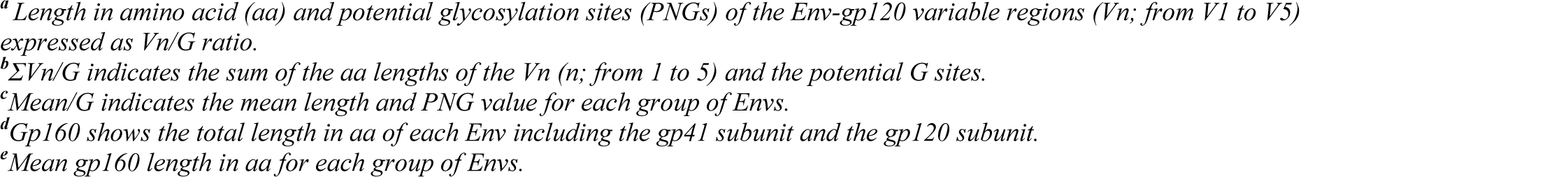
Molecular characteristics of HIV-1 Envs: sequence length and N potential glycosylation sites (PNGs) in the variable loops (Vn) of the gp120 subunit.

Regarding the PNGS in the sequences, many of the 24 relevant sites previously described (52–55) were present in these set of viral glycoproteins. However, major differences were observed in the aa extension of the loops with a progressive acquisition of more PNGS in the Modern Envs (**Table 2**). Glycan at N289 site was more present in LTNP-ECs, vLTNPs and Old viruses but is not present in Modern ones. Position N362 which is N proximal to the CD4 binding “DPE” motif (positions 368-370HXB2 sequence) was conserved in LTNP-EC, Viremic and Old but was only present in two of the Modern Envs. It is interesting to highlight that changes also occurred in the viral transmembrane gp41 protein in glycan N816 that was dominant in LTNPs but not in chronic individuals (Old and Modern).

It is interesting to mention that the trend in Env length increase follows the same pattern that the functional growth of the Env shown in the distinct viral characteristics (see **Figures 3 to 6**). We observed a good correlation between the genetic distance to the subtype B ancestor sequence obtained from Los Alamos National Laboratory HIV Database (LANL database, http://www.hiv.lanl.gov) and the functionality of viral Env proteins analysed (**Figure 9**). In general, the lower evolutionary sequences (less genetic distance to subtype B ancestor) are those with lower functionality (LTNP-ECs) and the higher evolutionary sequences are those with higher functionality (Moderns). In summary, the viral Envs with the most efficient characteristics are found within the Envs of the Modern group that also show the longer gp160 proteins, with more glycosylated sites and higher distance to the subtype B ancestor.

**Figure 9.**
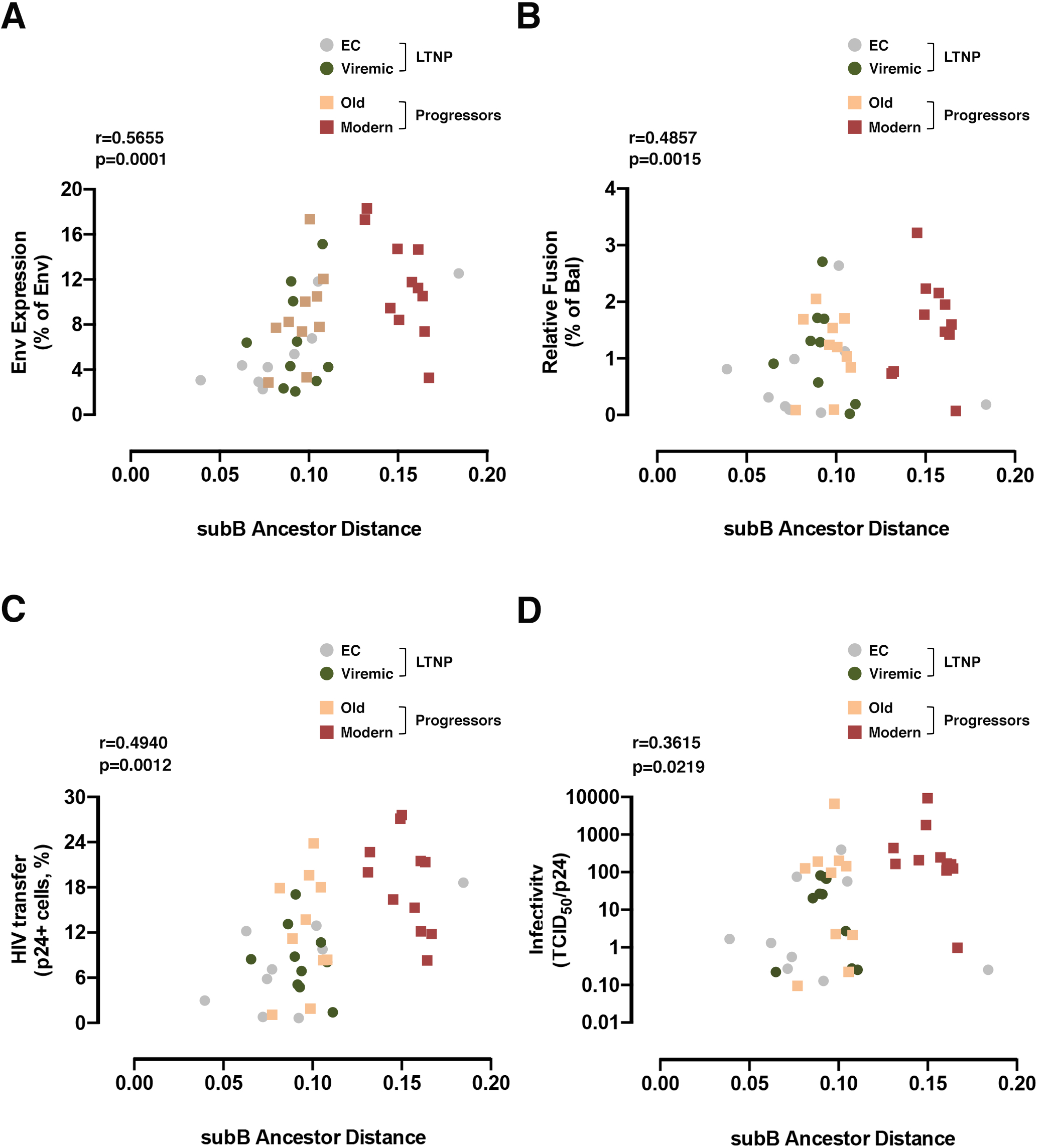
Correlation of the expression, fusion, transfer and viral infectivity Env characteristics with the nucleotide genetic distance to subtype B ancestor. Correlation between genetic distance to subtype B ancestor of all Envs of the different groups and Env expression (A), Relative fusion (B), HIV Transfer (C) and Infectivity (D). LTNP-ECs (*gray circle*), vLTNPs (*green circle*), Old patients (*orange square*) and Modern patients (*red square*). The correlations were calculated with a nonparametric Spearman test (p and r values are shown, *top left*). Values of Env expression, Relative fusion, HIV transfer and Infectivity are mean ± S.E.M. of three independent experiments.

## Discussion

HIV-1 infected individuals display a wide spectrum of clinical progression rates. The causes of this dispersion are multiple and associated with the operation of numerous combinations of host genetic, immunological and viral factors. In this work, we studied the potential contribution of viral Env glycoprotein characteristics to the clinical outcome of HIV-1 infection in HIV+ individuals with different clinical status.

The different groups of patients were defined by their clinical characteristics, distinct VLs and isolation dates because several studies have described a clear correlation between patients’ VL and the likelihood of virus transmission, disease progression and pathogenesis (56–63).

Although viral control in HIV-1 individuals has been linked to the host-immune responses (10, 64), other researchers and our group, however, stablished, in previous works, a direct connection between deficiencies in HIV-1 Env-associated functions and long-term viremia control in LTNP-ECs (17, 18, 20). The Envs from these LTNP-EC individuals were ineffective in the CD4 binding and in the subsequent functions: viral signaling, fusion and cell entry. These Env characteristics ensued in low replication and transmissibility of the virus (18, 19, 43, 45). All these data strongly support the role of the viral Env in the LTNP-EC phenotype and viral pathogenesis.

In the present work, we extended these observations to more Env from non-progressor subjects, which are not associated with a cluster of infection, in comparison to different sets of progressor chronic individuals. The Envs characteristics from LTNP individuals (EC and Viremic) were compared with those of individuals with progressive infection (Old and Modern). We investigated the defects in the association of Envs with the CD4, membrane fusion impairment and the cell-to-cell virus transfer and viral infection capacities. Viral Envs from LTNPs showed the lower binding capacity to the CD4 receptor and this initial inefficient Env/CD4 interaction led to a deficiency in membrane fusion and virus cell-to-cell transfer capabilities. The properties of the Env from LTNPs were not due to the ancestral origin of the LTNPs viruses isolated in the late 80’s and 90’s, because the chacteristics of the Old viruses which were contemporary to the LTNPs did not showed these limited functional characteristics. On the contrary, Envs from progressors (Old and Modern) presented efficient CD4-mediated viral functionality that triggered an effective membrane fusion and viral transfer. Thus, we disclosed that there is a clear correlation between the level of viral fusion, the transfer capacity of the viral Env and viral infectivity. The observed differences between the characteristics of the Envs from these groups could not be associated with viral tropism, because all the *env* nucleotide sequences from the studied viruses, showed an R5 tropism (Web PSSM, https://indra.mullins.microbiol.washington.edu/webpssm/). In summary, viral Envs from LTNPs exhibited non-functional characteristics (**Figures 3-6**) in comparison with those from viruses of the progressive infection groups, supporting the concept that the properties of the Envs were associated with viral control and the clinical progression rate of the HIV-1 individuals. In spite of the limited sampling, because of the difficult and laborious viral characterization of the viral phenotypes, we observed statistically significant differences between the characteristics of the Envs of viruses from LTNP-ECs and the Moderns. Also, if we consider the Env characteristics from all clinical groups, there is a consistent and recurrent tendency, although with no statistical power in some cases, to gain functionality in the viral Envs from the LTNP individuals (LTNP-ECs and vLTNPs), to those of the progressive groups (Old and Modern).

Remarkably, the increase in Env functionality also correlated with longer and more glycosylated proteins. The aa length and PNGs’ profile of the Envs from the individuals of the distinct clinical groups showed that the studied Envs tend to increase length and glycosylation over the course of the epidemic as previously described (see (49, 51)). We observed that Env changes accumulated essentially in the V1, V2, V4 and V5 loops, as previously shown in works relating the role of V1 and V4 loops in the CD4 binding and neutralization (65–68) and viral cell-to-cell transfer capacity (50, 69, 70). Regarding specific changes detected in our study, the loss of the N362 PNGs (position in the HXB2 isolate; group M, subtype B (HIV-1 M:B_HXB2R: NCBI:txid11706)) which was prevalent in the EC, Viremic and Old but not in the Modern Envs groups could be associated with the gain of functionality in the Envs. However, the opposite effect with more efficient fusion and transfer capacity was found in Australian viruses with the N362 glycosylation site (55). The potential role of the other changes in PNGs detected in our study need to be further investigated. Besides these important changes, it is clear that point mutations could have a significant impact in the viral characteristics and HIV pathogenesis (71, 72). The variants of concern (VOCs) of the pandemic severe acute respiratory syndrome coronavirus (SARS-CoV-2) unfortunately are reminding us (73, 74). Thus, the contribution of the individual mutations deserves further studies but it is now out of the scope of the present work.

In contrast with the more significative changes detected in the V2 and V5 loops, it is important to point the stability in length and glycosylation of the V3 loop. This structure is key for viral tropism (75–79) and for the correct CD4 Env binding as revealed with anti-V3 neutralizing antibodies that abrogate Env-CD4 interaction (80, 81).

In this study, we confirmed the inefficient functionality of the Envs from LTNP-EC individuals previously described for a cluster of viruses (18, 20), but extended to HIV+ individuals controlling viremia which are not clustered by the same transmitted/founder (T/F) virus. Also, a gain of Envs functionality from those of the LTNP individuals to the chronic not controlling individuals was identified. This improvement was detected in every Env characteristic analyzed; expression, fusion, virus transfer and infectivity. Interestingly, this functional growth of viral Env was associated in this study with length and PNGs increases in the variable loops. This increase was also reported in studies analyzing the susceptibility, neutralization sensitivity, co-receptor binding, host range and viral phenotype (49). This increase in the V1-V2 length and PNGs has also been detected thorough chronic infections from early to late viral Env sampling like in our work (49). Likewise in a group of individuals infected with closely related viruses higher PNGs density has been observed in the V1-V5 region of the gp120 during chronic infection compared to those oberved during the early acute infection phase (82). In viruses from the HIV-1 subtype B, it seems that early after viral transmission to a new host a selection for viral variants with shorter variable regions and a reduced degree of PNGs occurs (83). The growth in functionality of the viral characteristics was also correlated with the genetic distance of the sequences to the subtype B ancestor. Genetic variability in *env* gene has been associated with an increase in viral infectivity and replication capacity (84–89). These changes could facilitate viral replication by increasing viral fitness that favors the escape from the immune response and anti-retroviral therapy (ART) failure (90–99).

The non-functional characteristics of the primary Envs of LTNP individuals (ECs and Viremics) resulted in poor viral replication and very limited evolution that could allow the efficient immune control of HIV-1 infection and pathogenesis. It has been reported that in a LTNP-EC patient that followed discontinued ART, the V1 domain of his HIV-1 strain that retained good infectivity and replicative capacity included two additional N-glycosylation sites and was placed in the top 1% of lengths among the 6,112 Env sequences analyzed in the Los Alamos National Laboratory online database (100).

Therefore, it is conceivable that the functional characterization of the inefficient HIV-1 Envs could be significant in the development of a new generation of immunogens. Indeed, attenuated HIV or simian immunodeficiency virus (SIV) vaccines (LAHVs or LASVs) have been postulated as therapeutic vaccine strategies (101–107). However, further antigenic and immunogenicity work is needed to disclose the potential implications of these non-functional HIV Envs in the vaccine/cure field.

In summary, in this work, we exposed that the characteristics of the viral Envs from different groups of HIV-1 infected individuals could be associated with the short or long-term VL control and the clinical progression rate of the infection. The non-functional HIV-1 Envs could help in the development of new strategies for functional cure and virus eradication. Our data support the hypothesis that the functionality of viral Envs is a crucial characteristic for the control of viral infection, replication and pathogenesis.

## Material and methods

### Viral envelopes

Forty-one viral envelopes (Envs) were obtained from samples of different origins: the HIV HGM BioBank integrated in the Spanish AIDS Research Network (RIS-RETIC, ISCIII) (samples 1,2,3,6,7,8,13,14,15,16,17,18,19), the Centro Sanitario Sandoval, Hospital Clínico San Carlos (samples 21,22,24,28,30,31,32,33,36,37,38,39,40,42,43,44,45,46,49,50,51,52), the irsiCaixa Research Foundation (samples 9,10,11,12) and from Hospital Xeral de Vigo (samples 26,27). Samples were obtained in three different phases of the Spanish epidemic from 1993-94, 2004-2005 and 2013-2014. Samples were processed following current procedures and frozen immediately after their reception. All patients participating in the study gave their informed consent and protocols were approved by institutional ethical committees. Identification numbers and characteristics are found in **Table1**.

### Ethics Statement

Samples were obtained from participants who gave informed consent for genetic analysis studies and they were registered as sample collection in the Spanish National Registry of Biobanks for Biomedical Research with number C.0004030. The consents were approved by the Ethical and Investigation Committees of the “Centro Sanitario Sandoval” (Madrid) and the samples were encoded and de-identified in these Centers. All clinical investigations were conducted according to the principles expressed in the Declaration of Helsinki. The studies were approved by the Comité de Ética de la Investigación y de Bienestar Animal of the Instituto de Salud Carlos III with CEI PI 05_2010-v3 and CEI PI 09-2013 numbers.

### Generation of *env* gene expression plasmids

The *env* genes were amplified at limiting dilution by nested PCR from proviral DNA. The products were cloned into the pcDNA3.1D/V5-His’s Topo expression vector (Invitrogen) and NL4.3. The R5-tropic BaL.01-*env* (catalog number 11445) glycoprotein plasmid was from the NIH AIDS Research and Reference Reagent Program. Ten viral Envs were derived from 6 LTNP-EC patients, 10 clones from 6 Viremic LTNPs, 10 clones from 6 “Old” individuals (contemporary to LTNPs) and 11 clones from 10 recent “Modern” patients and NL4.3 and BaL.01 reference clones expression plasmids were transformed in DH5*α* cells, and clones sequenced to check the correct insertion of the *env* gene.

### Env expression and fusion assays

The Env expression plasmids were used to transfect HEK-293T cells with X-tremeGENE HP DNA Transfection Reagent (Sigma) in combination with either a Tat expression plasmid pTat for Env expression and fusion assays, or with the *env* defective HIV-1 backbone pSG3 plasmid for viral transfer assays (18, 19, 108). As a negative control, HEK-293T cells were transfected only with pTat and as a positive control we use the BaL and NL4.3 Envs. HEK-293T cells were chosen as effector cells since they provide sensitive measures of fusion even when using low fusogenic Env. 24 hours post-transfection, cells were collected, and tested for Env surface expression and also fusion activity. To test Env expression, 1x10^5^ Env/Tat co-transfected HEK-293T cells were incubated with 2G12 and IgGb12 monoclonal antibodies (mAbs; Polymun, Viena, Austria) at 6 μg/mL each for 45 minutes at RT. After washing the cells, the PE-labeled goat anti-human IgG (Jackson ImmunoResearch Laboratories) was added and incubated in the dark at room temperature for 15 minutes, as similarly reported (18, 19). Cells were washed, fixed in formaldehyde 1%, acquired in a Celesta flow cytometer (BD FACS Celesta) and analyzed using the Flow-Jo software (Tree Star Inc.) The percentage of Env-positive cells and the Mean Fluorescence Intensity (MFI) of these cells were used to evaluate Env expression.

To test fusion activity, 1x10^4^ Env/Tat-transfected or control Tat-transfected HEK-293T cells were mixed (ratio 1:1) in 96-well plates with CD4^+^CXCR4^+^CCR5^+^ TZM-bl reporter cells for 6 hours at 37°C. Luciferase activity was measured (Fluoroskan Accent, Labsystems) using Brite-Lite (PerkinElmer) and normalized to BaL-Env-mediated fusion. NL4.3 and BaL-Env expression plasmids were used as positive controls for Env staining and as reference value for fusion activity (BaL = 100%), as similarly reported (19, 108) (summarized in the scheme of **Figure 1B**).

### HIV-1 transfer/CD4 binding

To test viral transfer activity, which exclusively depends on the binding of gp120 to the CD4 molecule, Env expression plasmids were co-transfected with the Env-defective pSG3 plasmid in HEK-293T cells, as similarly reported (18, 19, 108). One day after transfection, 1x10^5^ HEK-293T cells were mixed at a 1:1 ratio in 96-well plates with primary CD4+ T lymphocytes freshly isolated from healthy donors by negative selection (CD4+ T-Cell Isolation Kit II, human, Miltenyi Biotec). Viral transfer was assessed after 24 hours of incubation at 37°C in permeabilized (FIX & PERM Cell Permeabilization kit, Invitrogen Life Technologies) and stained cells with the anti-HIV-1 p24 KC57 mAb (anti HIV core antigen RD1 labelled, IZASA) for 20 minutes in the dark at RT. Then, the cells were washed and fixed in formaldehyde 1%, and acquired in a Celesta flow cytometer (BD FACS Celesta) and the content of p24 in gated CD4+ T cells and gated HEK-293T cells was analyzed using the Flow-Jo software (Tree Star Inc.). The percentage of p24+ HEK-293T cells was used as a control for transfection efficiency and was similar among all experiments. Since co-receptor binding or fusion activity are not required for viral transfer, the frequency of p24+/CD4+ T cells was a direct measure of the amounts of HIV-1 virions bound to or taken up by target cells (summarized in the scheme of **Figure 1C**).

### Infectivity assay

Cloned viral Envs were used to generate pseudoviruses by co-transfection with pSG3 plasmid of HEK-293T cells as indicated above and tested in TZM-bl cells to determine the infectivity capacity. Serial Dilutions of the pseudoviruses generated with the different Envs of the different groups of patients were made in a 96-well plate. Then, 1x10^5^ TZM-bl cells were added to the pseudoviruses with DEAE dextran hydrochloride (Sigma) at 18 µg/mL. After 48 hours of incubation at 37°C, luciferase activity was measured (Fluoroskan Accent, Labsystems) using Brite-Lite (PerkinElmer). Uninfected TZM-bl cells were used as a negative control. The TCID_50_ (Median Tissue Culture Infectious Dose) value was calculated with Montefiori template and normalized with the viral concentrations (summarized in the scheme of **Figure 1D**).

### Phylogenetic Analysis

The evolutionary history was inferred by using the “maximum likelihood” (ML) method and JTT matrix-based model (109). The tree with the highest log likelihood (-49687,86) is shown. The percentage of trees in which the associated taxa clustered together is shown next to the branches. Initial tree(s) for the heuristic search was(were) obtained automatically by applying Neighbor-Join and BioNJ algorithms to a matrix of pairwise distances estimated using the JTT model, and then selecting the topology with superior log likelihood value. A discrete Gamma distribution was used to model evolutionary rate differences among sites (5 categories (+G, parameter = 0,6825)). The rate variation model allowed for some sites to be evolutionarily invariable ([+I], 18,05% sites). The tree is drawn to scale, with branch lengths measured in the number of substitutions per site. This analysis involved 140 aa sequences. All positions with less than 95% site coverage were eliminated (i.e., fewer than 5% alignment gaps), and missing data and ambiguous bases were allowed at any position (partial deletion option). There were a total of 829 positions in the final dataset. Evolutionary analyses were conducted in MEGA X (110).

Nucleotide sequences have been deposited in GeneBank under the following numbers: KC595156, KC595162, KC595225, KC595227, KC 595189, MH605987, MH605986, KC595190, MH605988, MH605992, MH605991, MH605970, MH605971, KC595223, KC595222, MH605972, MH605975, MH605976, MH605978, MH605973, MH605979, MH605980, MH605981, MH605982, MH605983, MH605984, MK394184, MK394185.

### Statistical analysis

Data and statistical analyses were performed using GraphPad Prism, version 6.07 (GraphPad Software). Significance when comparing groups was determined with a nonparametric Kruskal-Wallis or by nonparametric Dunn’s test for multiple comparisons. A nonparametric Spearman test was used to calculate correlations.

### Data Availability

All “accession numbers” and “data” of this work are available.

## Acknowledgements

We want to particularly acknowledge the patients in this study for their participation and to the HIV BioBank integrated in the Spanish AIDS Research Network and collaborating Centres (http://hivhgmbiobank.com/donor-area/hospitals-and-centres-transferring-samples/?lang=en) for the generous gifts of clinical samples used in this work. The HIV BioBank, integrated in the Spanish AIDS Research Network, is partially funded by the RD16/0025/0019 project as part of the Plan Nacional R+D+I and cofinanced by ISCIII-Subdirección General de Evaluación and el Fondo Europeo de Desarrollo Regional (FEDER). The clinical follow-up of Drs. Carmen Rodriguez, Mar Vera and Jorge Del Romero (Centro Sanitario Sandoval), Eulalia Grau (Hospital Germans, Trias y Pujol; irsiCaixa, Badalona) is greatly appreciated.

